## Supplementary information for "Identification of picornavirus proteins that inhibit *de novo* nucleotide synthesis during infection"

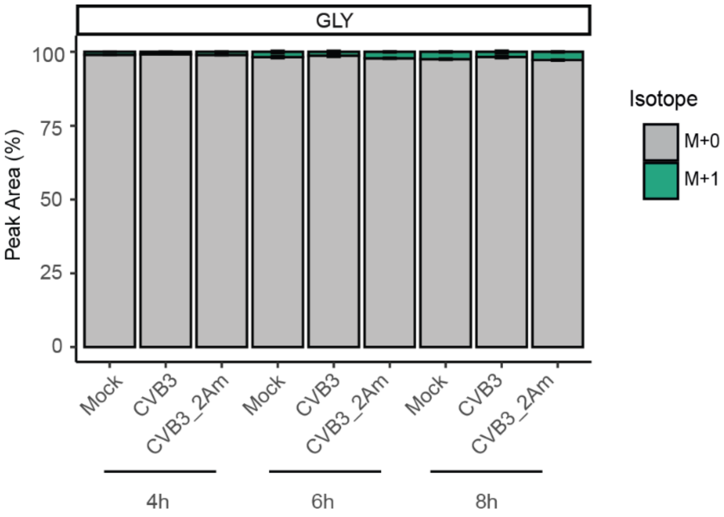


**S1. Glycine is marginally labeled by $^{15}N_2$-glutamine.** $^{15}N_2$-glutamine isotope tracing study in mock- , CVB3-2Am- and CVB3-infected HeLa R19 cells (MOI 5 TCID50/ml, three replicates treatment and per isotope). Cells were infected, lysed at 4,6 or 8 hpi and measured by LC-MS to identify metabolites and quantify the different isotopologues. The labeling of glycine by $^{15}N_2$-glutamine is shown.

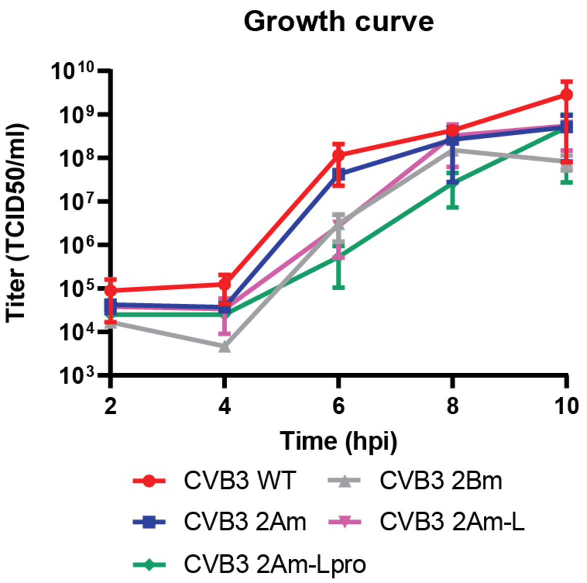


**S2. Growth kinetics and characteristics of the recombinant CVB3 viruses.** A) Growth kinetics of the recombinant CVB3 viruses in HeLa R19 cells, titrated on HeLa R19 cells (mean and SD of triplicates; one experiment, MOI 5 TCID50/ml). The cells were infected, lysed at 2, 4, 6, 8 and 10 and titrated (on HeLa R19 cells) to determine the TCID50/ml.

**A**

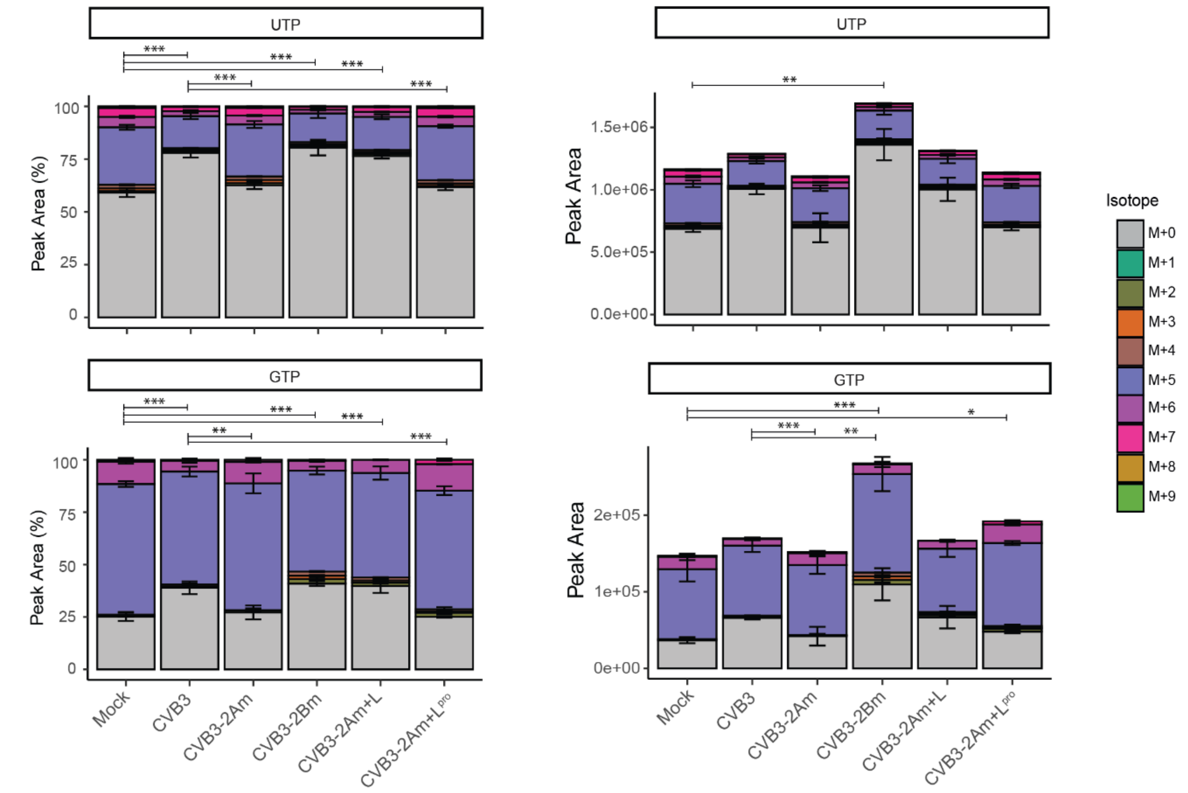


**B**

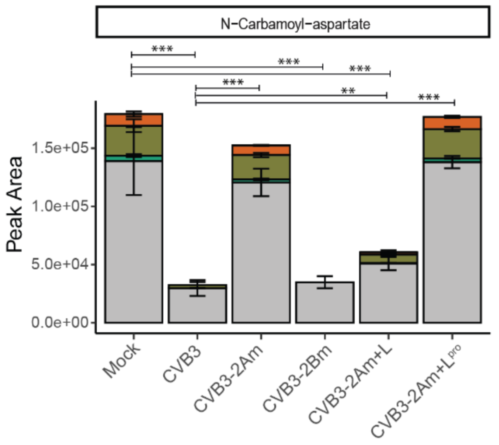


**S3. 2A$^{pro}$ and L restrict de novo nucleotide synthesis, while L$^{pro}$ does not.** $^{13}C_6$-glucose isotope tracing study in mock- and CVB3-, CVB3-2Am, CVB3-2Am+L, CVB3-2Am+L$^{pro}$ infected HeLa R19 cells (MOI 5 TCID50/ml, three replicates). Cells were infected, lysed at 6 or 8 hpi and measured by LC-MS to identify metabolites and quantify the different isotopologues. Data obtained at 6 hpi are shown here. A) Peak areas and isotopologue distribution of UTP and GTP at 6 hpi. D) Peak areas of N-carbamoyl-aspartate at 6 hpi. For statistical analysis, linear mixed effect models with an interaction of time and treatment and a random effect of replicate were performed. A rank transformation on the data was performed to ensure a normal distribution of the residuals. For GTP, a normal distribution of the residuals could not be assumed for the fractional data and therefore a non-parametric linear mixed effect model with an interaction of time and treatment and a random effect of replicate was performed. p-values between specific groups were calculated by performing a contrast analysis, in which total labeled fraction (A, left panels) or total peak areas (A, right panels & B) were compared between groups. *p < 0.05, **p < 0.01, ***p < 0.001.

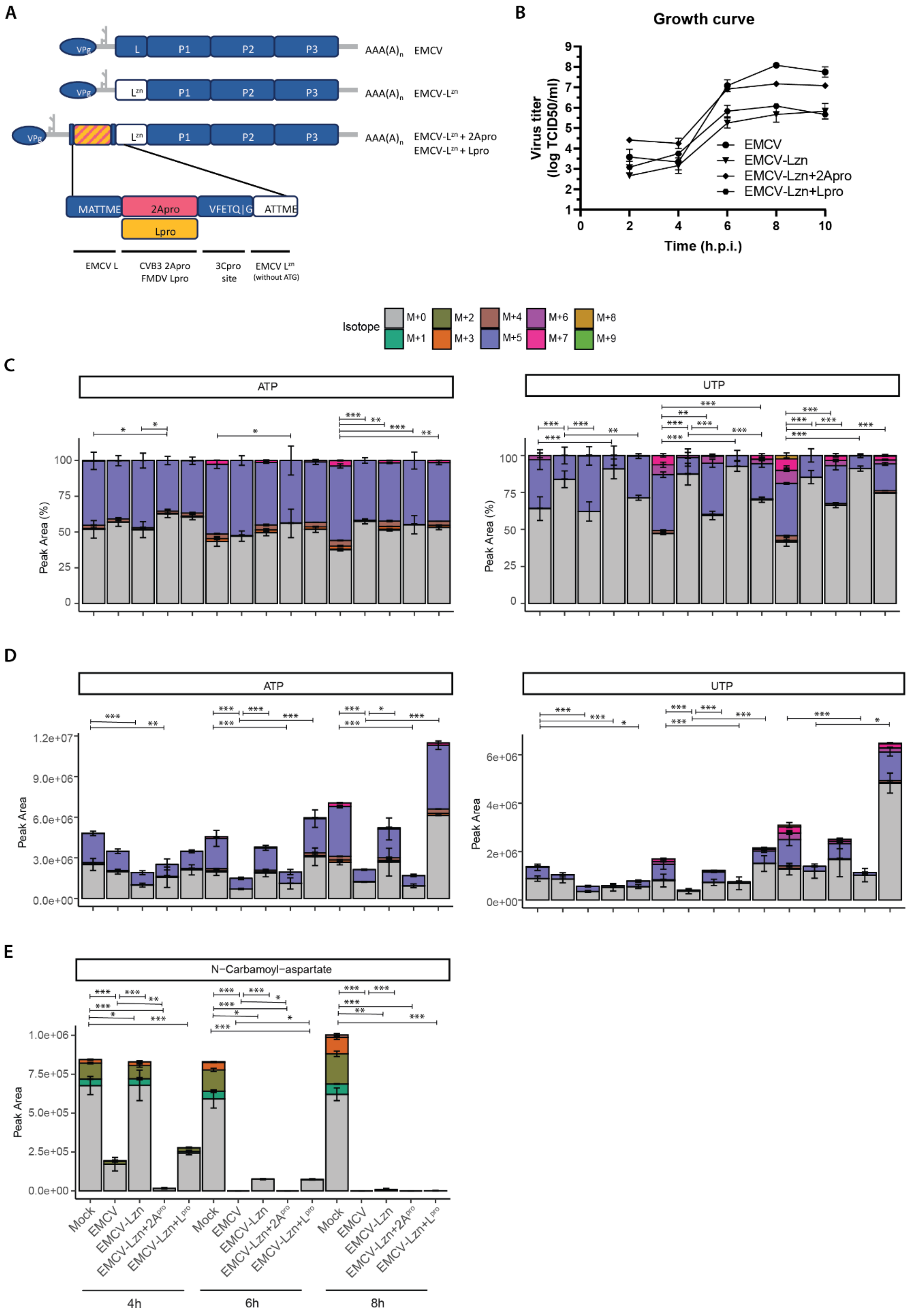


**S4. $2A^{pro}$ and L restrict *de novo* nucleotide synthesis, while $L^{pro}$ does not.** $^{13}C_6$-glucose isotope tracing study in mock- and EMCV-, EMCV-$L^{zn}$-, EMCV-$L^{zn}$+$2A^{pro}$- and EMCV-$L^{zn}$+$L^{pro}$-infected HeLa R19 cells (MOI 5 TCID50/ml, three replicates; the mock and EMCV conditions of this experiment have already been shown in our previous work (34)). Cells were infected, lysed at 4,6 or 8 hpi and measured by LC-MS to identify metabolites and quantify the different isotopologues. two samples were removed from analysis (one replicate of 8 hpi EMCV and one replicate of 8 hpi EMCV-$L^{zn}$+$L^{pro}$), because of a technical defect. A) Schematic representation of EMCV and the EMCV recombinant viruses that were used in this study. B) Growth curve of EMCV and the recombinant EMCV viruses in HeLa R19 cells. C) Absolute peak areas and isotopologue distribution of UTP and ATP. D) Absolute peak areas of N-carbamoyl-aspartate. For statistical analysis, linear mixed effect models with an interaction of time and treatment and a random effect of replicate were performed. For D) A rank transformation on the data was performed to ensure a normal distribution of the residuals. For N-carbamoyl-aspartate in E), a normal distribution of the residuals could not be assumed and therefore a non-parametric linear mixed effect model with an interaction of time and treatment and a random effect of replicate was performed. p-values between specific groups were calculated by performing a contrast analysis, in which total labeled fraction (C) or total peak areas (D & E) were compared between groups. *$p < 0.05$, **$p < 0.01$, ***$p < 0.001$.

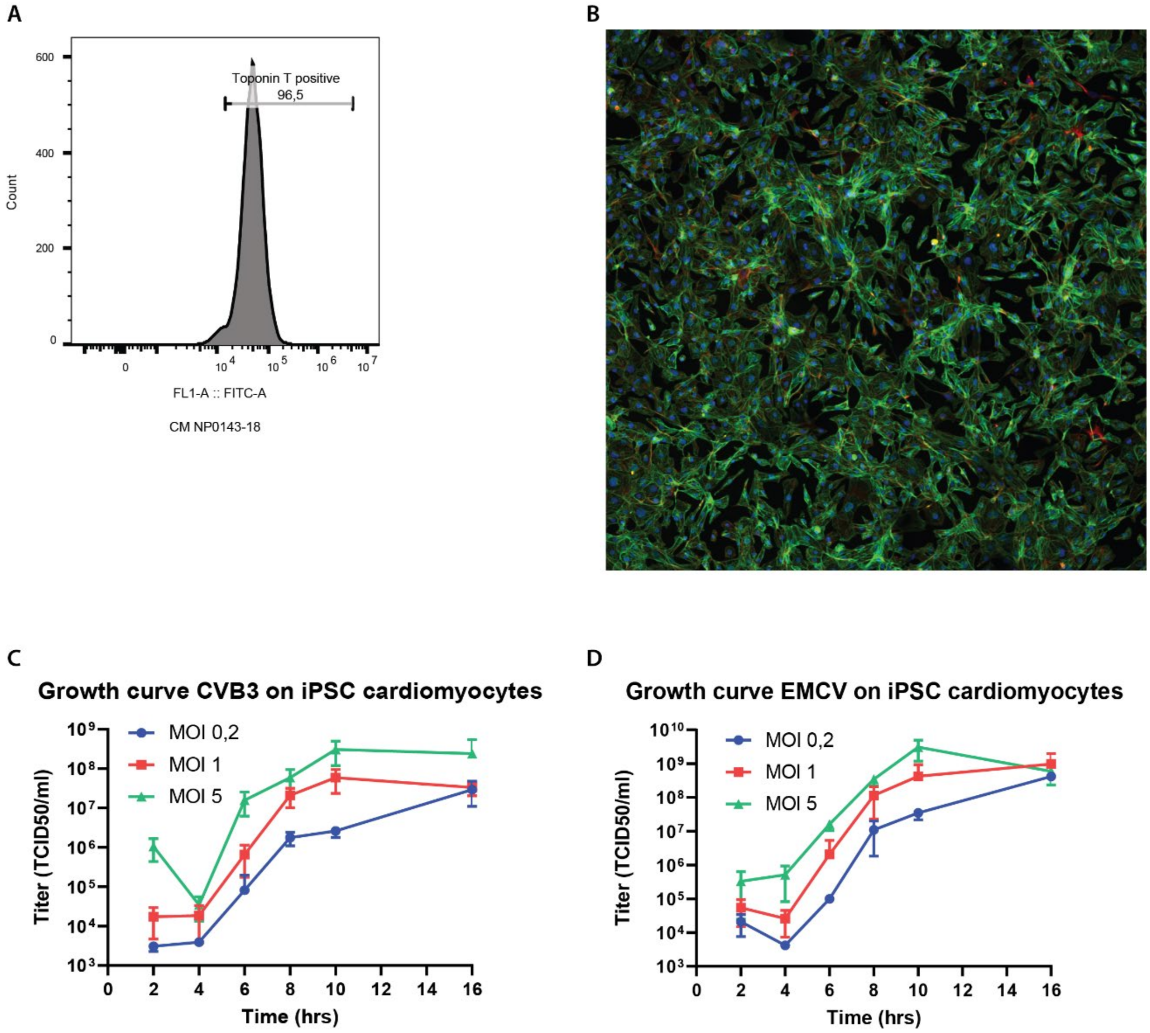


**S5. Growth kinetics CVB3 and EMCV in hiPSC-CMs.** A) Representative flow cytometry plot of Troponin T positive hiPSC-CMs (day 13), showing highly pure differentiations (96.5% Troponin T+). B) Representative fluorescent immunostaining of Troponin T positive hiPSC-CMs (in green) and Vimentin positive non-CMs (in red) with hoechst counterstained nuclei (in blue), showing normal hiPSC-CM sarcomere structure and high purity of differentiation. C), D) Growth kinetics of CVB3 (C) and EMCV (D) in hiPSC-CMs, titrated on HeLa R19 cells (mean and SD of triplicates; one experiment, MOIs (TCID50/ml) are calculated using HeLa R19 titers). The cells were infected, lysed at 2, 4, 6, 8, 10 and 16 hpi and titrated to determine the TCID50/ml.

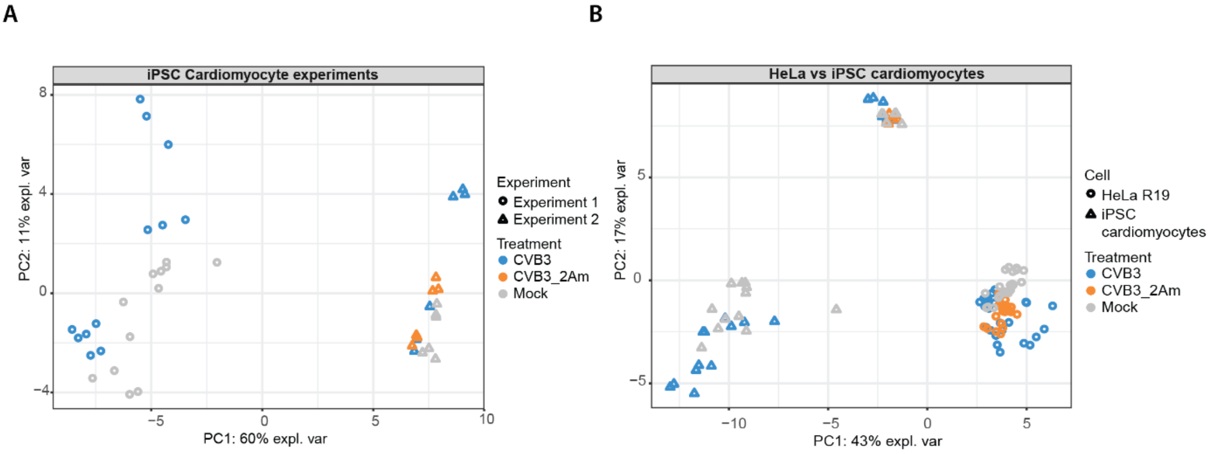

**S6. Variation in metabolic rewiring during CVB3 infection in hiPSC-CMs and compared to HeLa R19 cells.** $^{13}C_6$-glucose isotope tracing studies in mock-, CVB3- or CVB3-2Am infected hiPSC-CMs (MOI 5 TCID50/ml calculated from the HeLa R19 titer, three replicates) or HeLa R19 cells (MOI 5 TCID50/ml, three replicates). Cells were infected, lysed at 2.5, 5, 7.5, 10 hpi (hiPSC-CMs) or lysed at 2, 4, 6, 8, 10 hpi (HeLa R19 cells) and measured by LC-MS to identify metabolites and quantify the different isotopologues. A) PCA plot of the experiments with hiPSC-CMs. B) PCA plot of the experiments of both hiPSC-CMs and HeLa R19 cells.

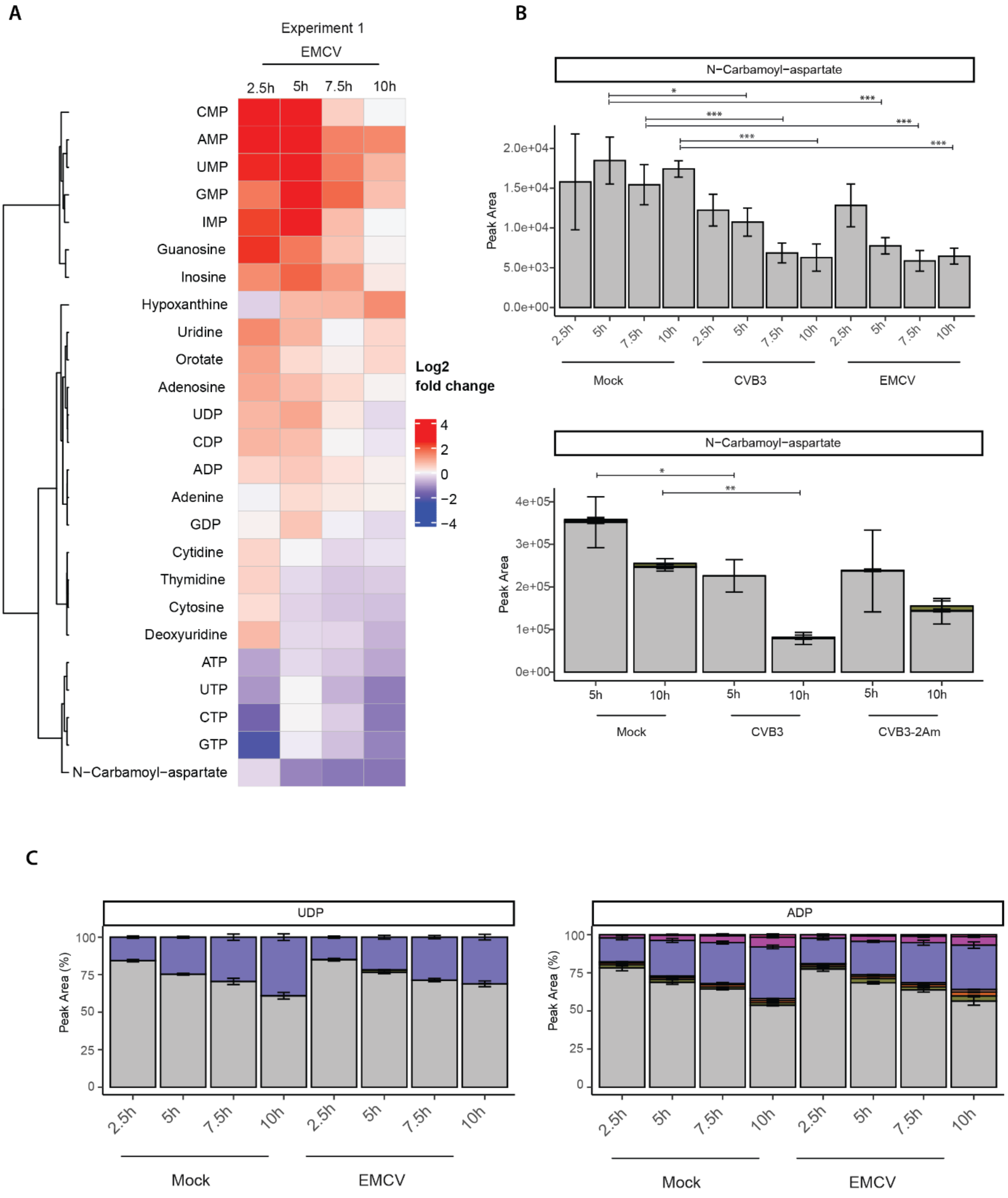

**S7. CVB3 and EMCV elevate levels of purine and pyrimidine metabolites in hiPSC-CMs.** $^{13}C_6$-glucose isotope tracing study in mock-, CVB3-, CVB3-2Am or EMCV infected hiPSC-CMs (MOI 5 TCID50/ml calculated from the HeLa R19 titer, three replicates). Cells were infected, lysed at 2.5, 5, 7.5, 10 hpi and measured by LC-MS to identify metabolites and quantify the different isotopologues. A) Heatmap showing log2 fold changes of the purine and pyrimidine metabolites between EMCV infected- and mock-infected cells. Log2 fold changes are calculated based on the mean of three replicates. B) Absolute levels of N-carbamoyl-aspartate. C) Isotopologue distribution of UDP and ADP. To statistically analyze the data, linear mixed effect models with an interaction of time and treatment and a random effect of replicate were performed. A rank transformation on the data was performed to ensure a normal distribution of the residuals. p-values between specific groups were calculated by performing a contrast analysis, in which total peak areas (B) or total labeled fraction (C) were compared between groups. *p < 0.05, **p < 0.01, ***p < 0.001.

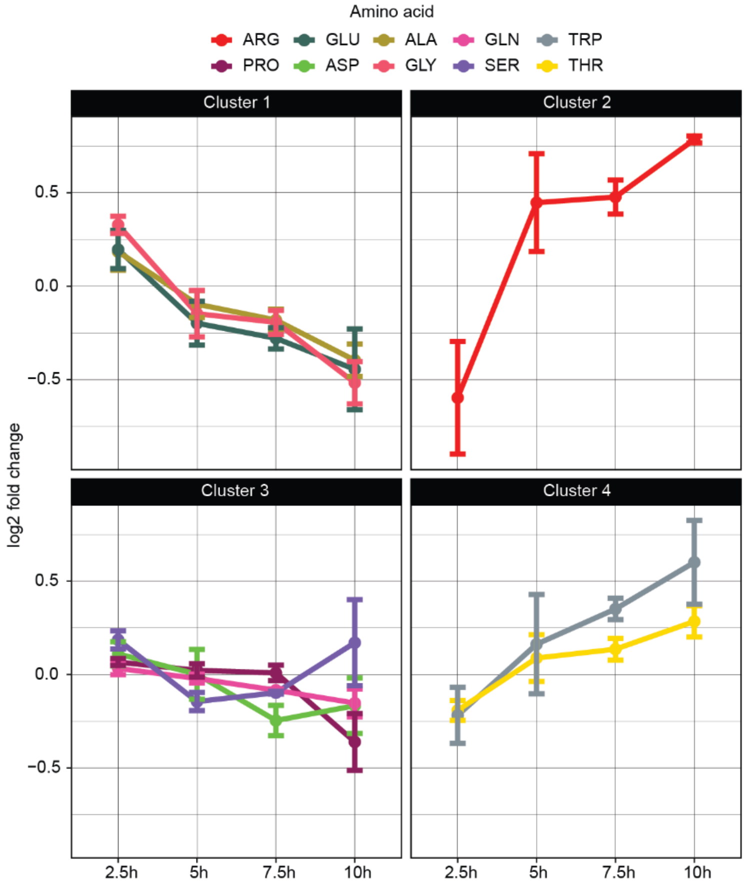


**S8. CVB3 and EMCV infection alters amino acid levels in hiPSC-CMs.** $^{13}C_6$-glucose isotope tracing studies in mock- or EMCV-infected hiPSC-CMs (MOI 5 TCID50/ml, three replicates per experiment). Cells were infected, lysed at 2.5, 5, 7.5 or 10 hpi and measured by LC-MS to identify metabolites and quantify the different isotopologues. The isotopologue distribution is not shown in this figure. A) Log2 fold changes of amino acid levels during EMCV infection over time.

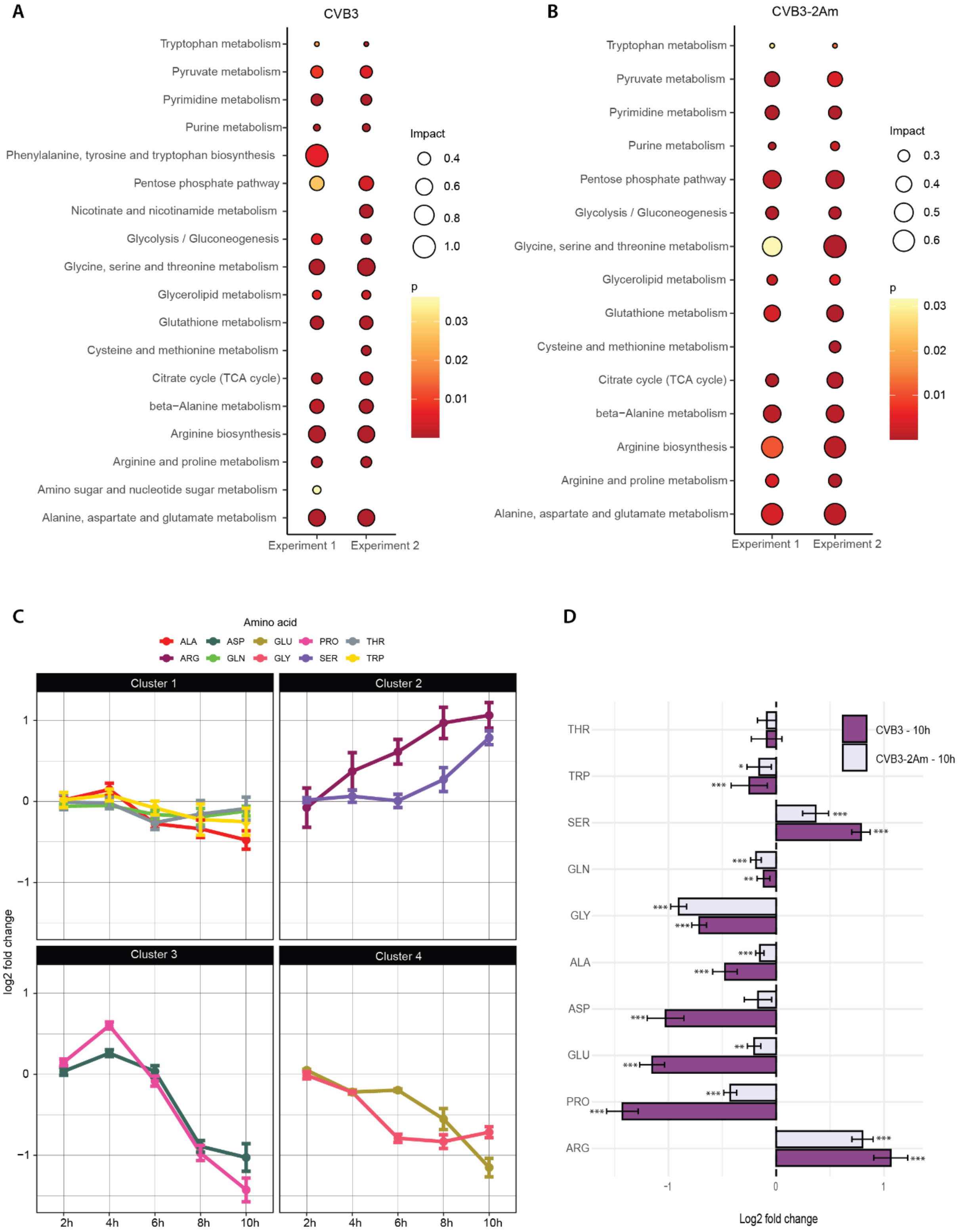


**S9. CVB3 modulates arginine, proline, aspartate and glutamate metabolism in HeLa R19 cells.** $^{13}C_6$-glucose isotope tracing studies in mock-, CVB3- and CVB3-2Am HeLa R19 cells (MOI 5 TCID50/ml, three replicates per experiment). Cells were infected, lysed at 2, 4, 6, 8 or 10 hpi and measured by LC-MS to identify metabolites and quantify the different isotopologues. The different isotopologues are not distinguished in this Figure. A) MetaboAnalyst pathway analysis of the two different metabolomic experiments performed on HeLa R19 cells infected with CVB3 at 8 hpi. B) MetaboAnalyst pathway analysis of the two different metabolomic experiments performed on HeLa R19 cells infected with CVB3-2Am at 8 hpi. C) Representative Log2 fold changes of amino acid levels during CVB3 infection over time (from experiment 1). D) Representative Log2 fold changes between CVB3-2Am- vs mock-infected HeLa R19 cells at 8 hpi of amino acids (from experiment 1). To statistically analyze the data, linear mixed effect models with an interaction of time and treatment and a random effect of replicate were performed. p-values between specific groups were calculated by performing a contrast analysis, in which total peak areas were compared between groups. *$p < 0.05$, **$p < 0.01$, ***$p < 0.001$.
